# Modeling Endothelial Dysfunction in Idiopathic Pulmonary Fibrosis: Bridging Mechanistic Insights and Translational Applications

**DOI:** 10.1101/2025.09.29.679236

**Authors:** Annalena Branz, Irina Shalashova, Anastasia M. Funk, Tarek Gensheimer, Agnieszka Wytyk, Susanne Wespel, Jochen Drescher, Christoph H. Mayr, Lavinia Neubert, Jan C. Kamp, Mark Kühnel, Danny Jonigk, Manfred Frick, Muriel Lizé

## Abstract

The alveolus, the lung’s primary gas exchange unit, relies on tightly coordinated interactions between epithelial and endothelial layers. In idiopathic pulmonary fibrosis (IPF), a progressive interstitial lung disease, this architecture is profoundly disrupted. While epithelial and mesenchymal compartments have been extensively studied, the role of pulmonary microvascular endothelial cells (PMVECs) in IPF pathogenesis remains underexplored.

Here, we characterize PMVEC alterations in IPF using single-cell RNA sequencing and spatial transcriptomics, identifying subtype-specific markers and demonstrating their progressive loss in fibrotic lungs. To model endothelial dysfunction, we established robust protocols for isolating and culturing primary human ECs and applied a pharmacologically relevant cytokine cocktail (IPF-RC) that mimics the IPF microenvironment. IPF-RC exposure induced hallmark features of endothelial injury, including VE-cadherin loss, increased ICAM1/VCAM1 signaling, impaired barrier integrity, and reduced wound healing and angiogenic capacity.

To address the need for translational tools in drug discovery, we optimized and validated a suite of functional, scalable test systems and their endpoints using both primary and commercial endothelial cells. These mechanistic assays reliably recapitulate fibrotic endothelial injury and enable quantitative assessment of therapeutic interventions. Notably, treatment with a cAMP analog partially restored endothelial function, supporting the utility of these models for regenerative and pharmacological screening.

Our findings position PMVECs as active participants in IPF progression and present novel, scalable test systems that bridge mechanistic insight with translational application. These models offer a valuable platform for identifying endothelial-targeted therapies aimed at restoring alveolar capillary integrity in fibrotic lung disease.

## 2. Introduction

Idiopathic pulmonary fibrosis (IPF) is a progressive and degenerative lung disease with limited therapeutic options **[1]**. Despite recent therapeutic advances, current treatments offer only modest benefits, and there remains an urgent need for disease-modifying strategies.

IPF pathogenesis involves a complex interplay between alveolar niche cells and systemic factors. The prevailing hypothesis suggests that recurrent injury to alveolar epithelial cells leads to metabolic dysfunction, abnormal activation, senescence, and impaired repair, ultimately distorting lung architecture **[2–5]**. Additionally, dysregulated epithelial cells can affect surrounding cell types, causing their activation and malfunctioning **[2,3,6]**. While epithelial and mesenchymal compartments have been extensively studied, the contribution of pulmonary microvascular endothelial cells (PMVECs) to disease progression remains poorly understood.

Endothelial cells (ECs) are essential for maintaining vascular homeostasis, regulating immune cell trafficking, and supporting alveolar integrity **[6]**. Despite their importance, ECs have received considerably less attention than epithelial and mesenchymal cells in IPF research. Emerging evidence suggests that ECs, like epithelial cells, may undergo repeated injury-repair cycles in response to aberrant signaling and metabolic stress **[7,8]**, potentially leading to altered cell fate and contributing to chronic inflammation through paracrine and juxtracrine mechanisms **[9,10]**.

Recent advances in single-cell transcriptomics have identified two distinct pulmonary microvascular EC subtypes-general capillary cells (gCaps) and aerocytes (aCaps) - which mirror the AT2-AT1 epithelial lineage relationship **[6,11]**. gCaps regulate vascular tone and immune responses and serve as progenitors for aCaps, which mediate gas exchange and leukocyte trafficking. While epithelial cell fate in fibrosis is well characterized, the behavior and fate of these EC subtypes during lung injury remain poorly understood. We hypothesize that fibrotic remodeling and devascularization in IPF lead to a depletion of gCaps and aCaps, contributing to disease progression.

Spatial transcriptomics and histopathological analyses have revealed aberrant endothelial niches in IPF lungs, including ectopic ECs with phenotypes reminiscent of basaloid epithelial cells **[12]**. These findings suggest that ECs may acquire disease-specific traits and actively contribute to fibrotic remodeling. Despite these insights, a major limitation in IPF research is the lack of robust, disease-relevant *in vitro* models to study endothelial dysfunction under fibrotic conditions. Such systems are essential for dissecting disease mechanisms, identifying therapeutic targets, and evaluating candidate compounds aimed at restoring microvascular function. Existing platforms - such as precision-cut lung slices, organ-on-chip systems, and cytokine cocktails - offer valuable mechanistic insights but are limited by poor endpoint characterization, biological variability, scalability, and throughput, restricting their utility in drug discovery.

To address this gap, we developed and validated pharmacologically relevant test systems that model fibrotic endothelial injury using both primary and commercial ECs. These assays enable quantitative assessment of endothelial dysfunction and therapeutic response, offering translational value for regenerative strategies and drug screening.

In summary, we characterize PMVEC alterations in IPF, define subtype-specific markers, and establish functional assays that recapitulate key features of endothelial injury. Our findings support the hypothesis that ECs are active contributors to IPF pathogenesis and provide a scalable platform for identifying endothelial-targeted therapies aimed at restoring alveolar capillary integrity.

## 3. Results

Recent data sets have begun to reveal alterations in ECs in IPF, particularly in PMVECs. However, the extent and nature of these changes remain insufficiently characterized. To support the identification of biomarkers with diagnostic and mechanistic relevance, we analyzed cell type–specific expression of selected endothelial marker genes using publicly available single-cell RNA sequencing (scRNA-seq) data from the endothelial compartment of IPF and control lungs (Fig. 1A) [12].

**Figure 1.**
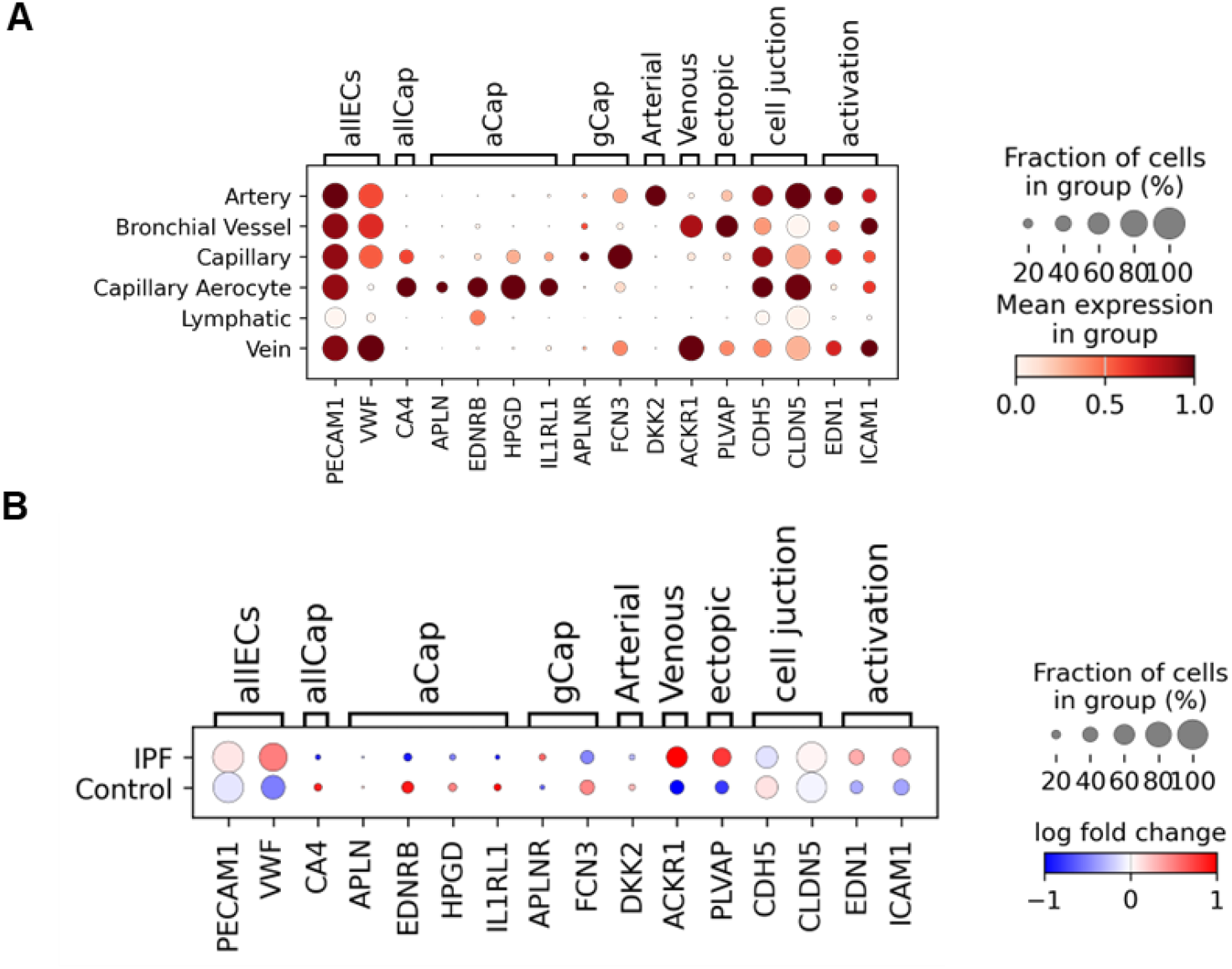
Single-cell RNA-seq reveals altered endothelial marker expression in IPF lungs. (A) Dot plots representing scRNA-seq data, visualizing the scaled expression of the indicated genes across various cell types. (B) Dot plots generated from scRNA-seq data, showing log fold-change in gene expression between IPF/control and control/IPF groups within the endothelial compartment. Data set from Mayr et al., 2024[12].

To investigate disease-associated changes, we compared fold changes in gene expression between IPF and control samples. Markers associated with aerocytes (aCap) - a specialized EC subtype involved in gas exchange - such as EDNRB, HPGD, and IL1RL1, were significantly downregulated in IPF. In contrast, markers indicative of endothelial activation (EDN1, ICAM1) and disease-induced ectopic ECs (PLVAP) were upregulated (Fig. 1B). Recognizing the limitations of scRNA-seq, particularly the isolation bias affecting fragile and elongated capillary cells, we validated these findings using spatial transcriptomics from formalin-fixed paraffin-embedded lung tissue sections. In control lungs, aCap and general capillary (gCap) markers were enriched in the alveolar niche, whereas IPF samples showed loss of microvascular markers and increased expression of activation markers in the same regions (Fig. A1A).

To further confirm endothelial alterations in IPF, we performed quantitative PCR on lung tissue from five control and six IPF patients. The results revealed significant downregulation of aCap markers (HPGD, EDNRB, IL1RL1) and gCap markers (FCN3, CA4) in IPF samples, while PLVAP expression remained unchanged (Fig. 2A). To ensure these differences originated from ECs, we isolated CD31^+^ cells from one control and one IPF donor and analyzed them at passage 5 following FACS enrichment. The expression profile mirrored tissue-level findings, with reduced aCap and gCap marker expression in IPF-derived ECs (Fig. A1B). Notably, we observed giant HPGD^+^, CD31^+^ cells measuring 200 µm or larger, consistent with early, immature aerocyte morphology adaptation described in literature (Fig. 2C). However, prolonged 2D culture led to progressive loss of microvascular markers and disappearance of these giant cells (Fig. A1C), highlighting the limitations of conventional culture systems.

**Figure 2.**
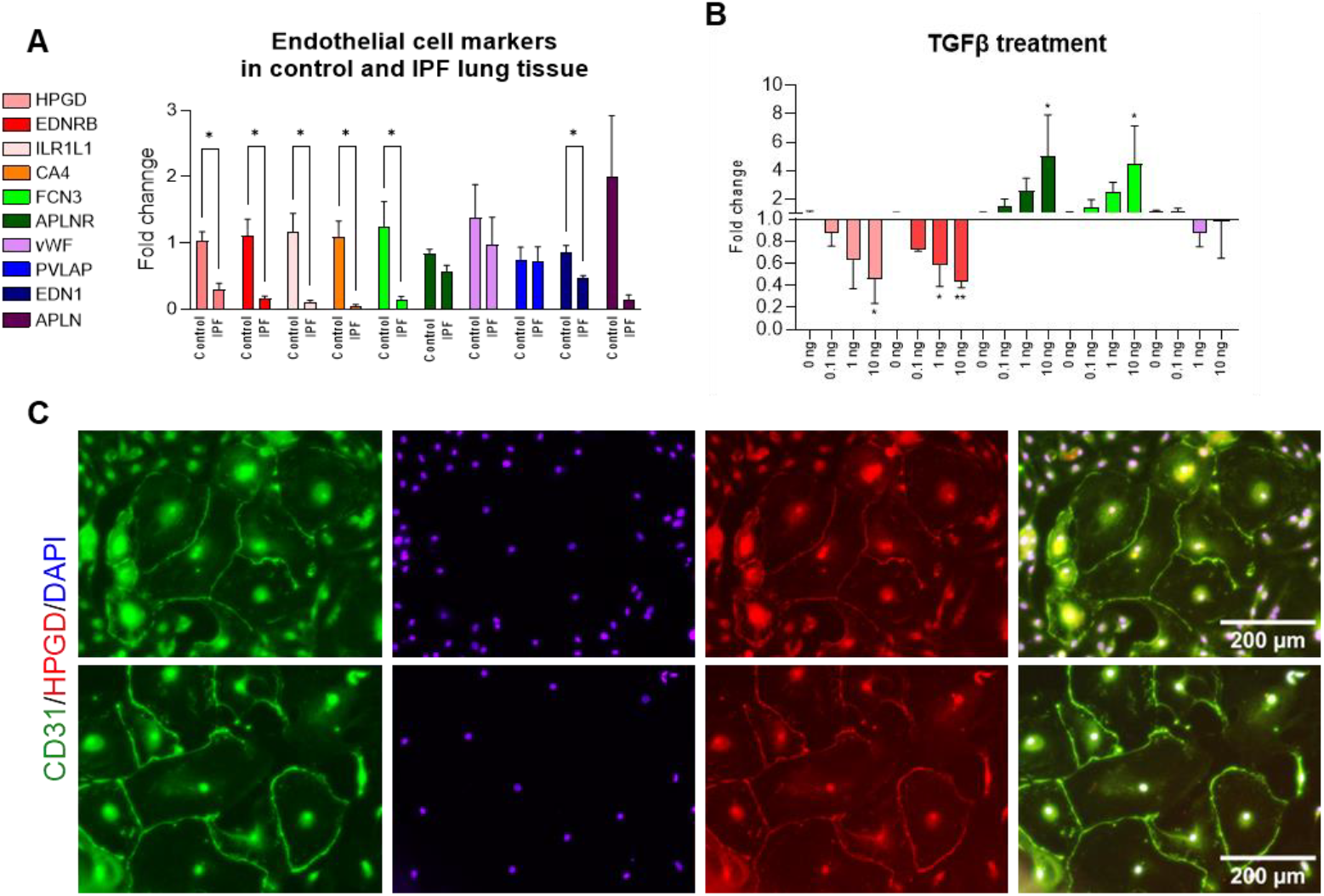
Validation of endothelial marker dysregulation in *ex vivo* lung tissue. (A) Expression of endothelial markers in control lungs (n = 5) and IPF lungs (n = 6) analyzed by real-time PCR. Statistics: unpaired t-test with Welch correction; *p* ≤ 0.05. Only significant results are labeled. (B) Expression of endothelial markers in precision-cut lung slices (PCLS) treated with TGFβ (0, 0.1, 1, 10 ng/mL) for 48 h. Statistics: one-way ANOVA with Dunnett’s multiple comparison test. Data shown as fold change relative to untreated control. Error bars: SEM from three donors (n = 3). Significance: *p* ≤ *0*.*05 (*), p* ≤ *0*.*01 (**), p* ≤ *0*.*001 (***), p* ≤ *0*.*0001 (****)*. (C) Immunofluorescence (IF) images of CD31^+^ primary cells at passage 0, days 10–12 post-seeding. CD31 (green), HPGD (red), DAPI (blue).

To better understand EC behavior under fibrotic conditions at early fibrotic stage, we employed human PCLS treated with TGFβ1 (Fig. 2B). A dose-dependent reduction in aCap markers (HPGD, EDNRB) was observed after 48 hours, while gCap markers (APLNR, FCN3) increased. Interestingly, vWF expression remained stable, suggesting selective modulation of PMVEC subtypes. It is important to note that PCLS contain multiple cell types, which may contribute to the observed transcriptional changes. Therefore, further validation using purified EC populations is warranted to confirm these findings.

The observation that endothelial cell subtypes undergo changes in fibrotic environments prompted us to investigate the fate and functional behavior of ECs derived from IPF versus control lung tissue. To this end, we isolated primary CD31^+^ endothelial cells from both patient groups and assessed their response to a disease-relevant cytokine cocktail (IPF-RC) [18], designed to mimic the inflammatory milieu of IPF lungs.

Endothelial cells play a critical role in maintaining the alveolar-capillary barrier required for efficient gas exchange. This barrier relies on tight junctions to prevent fluid leakage and regulate vascular permeability. Under pathological conditions, such as fibrosis, this integrity is compromised, leading to increased permeability and tissue damage. Although phenotypic differences between control- and IPF-derived ECs were subtle, IPF-derived cells exhibited signs of weakened junctional integrity, as evidenced by reduced VE-cadherin and CD31 expression (Fig. 3A–D). Both cell types responded to IPF-RC stimulation with increased expression of the monocyte adhesion molecule ICAM1 (Fig. 3A, B), indicating endothelial activation. However, no significant differences were observed between control and IPF-derived cells in tube formation assays (Fig. A2A). To further characterize endothelial activation, we quantified soluble biomarkers sICAM1 and sVCAM1 following IPF-RC treatment (Fig. 4A, B). Both control and IPF-derived ECs showed comparable responses, suggesting that prolonged culture and expansion in 2D conditions may lead to phenotypic convergence and loss of disease-specific traits (Fig. A1C).

**Figure 3.**
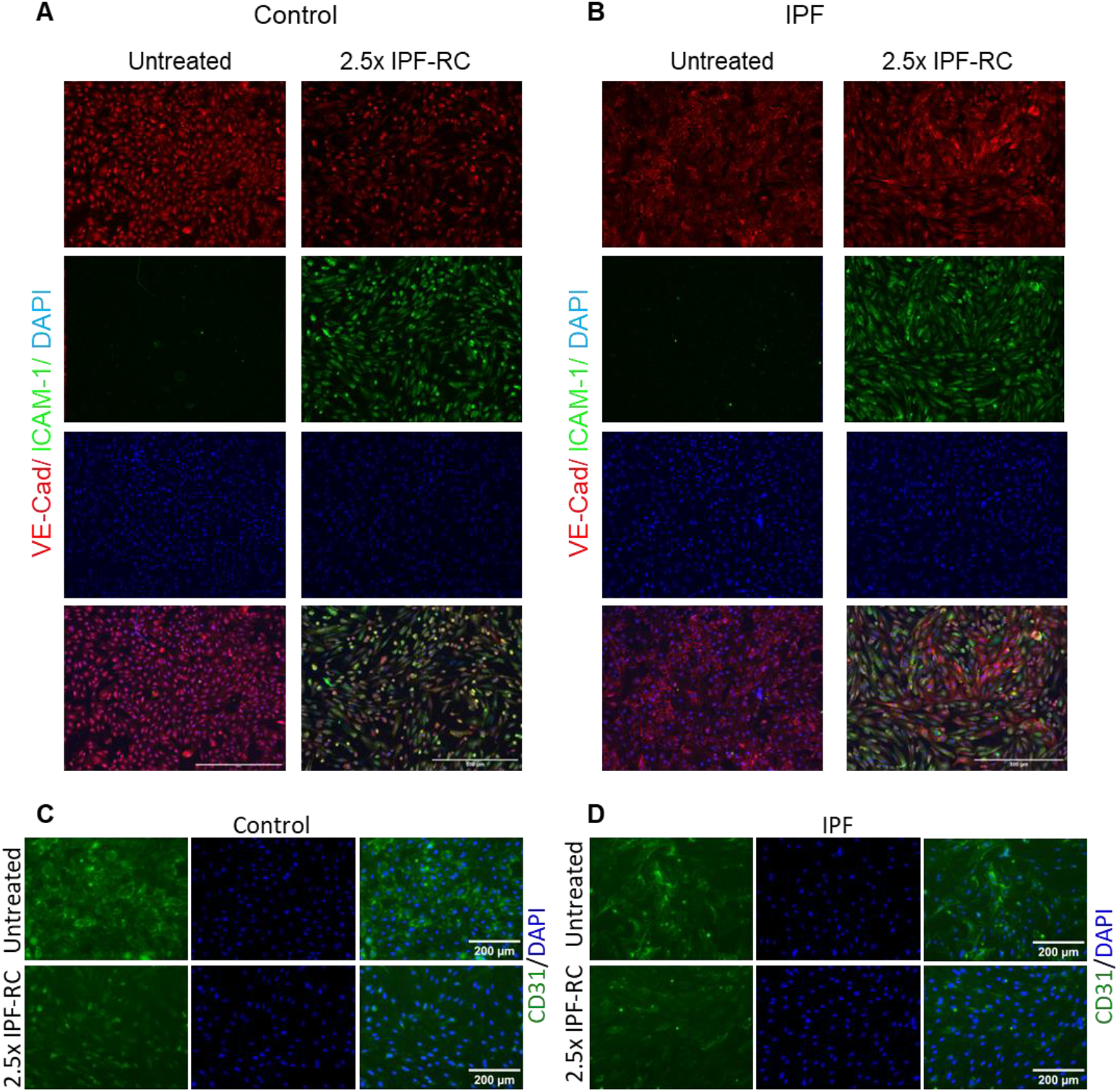
Confirmation of EC activation under IPF-RC stimulation. (A–B) Representative IF images of CD31^+^ cells from control (A) and IPF lungs (B) treated with 2.5× IPF-RC for 24 h. VE-cadherin (red), ICAM1 (green), DAPI (blue). (C–D) Representative IF images of CD31^+^ cells from control and IPF lungs treated with 2.5× IPF-RC for 24 h. CD31 (green), DAPI (blue).

**Figure 4.**
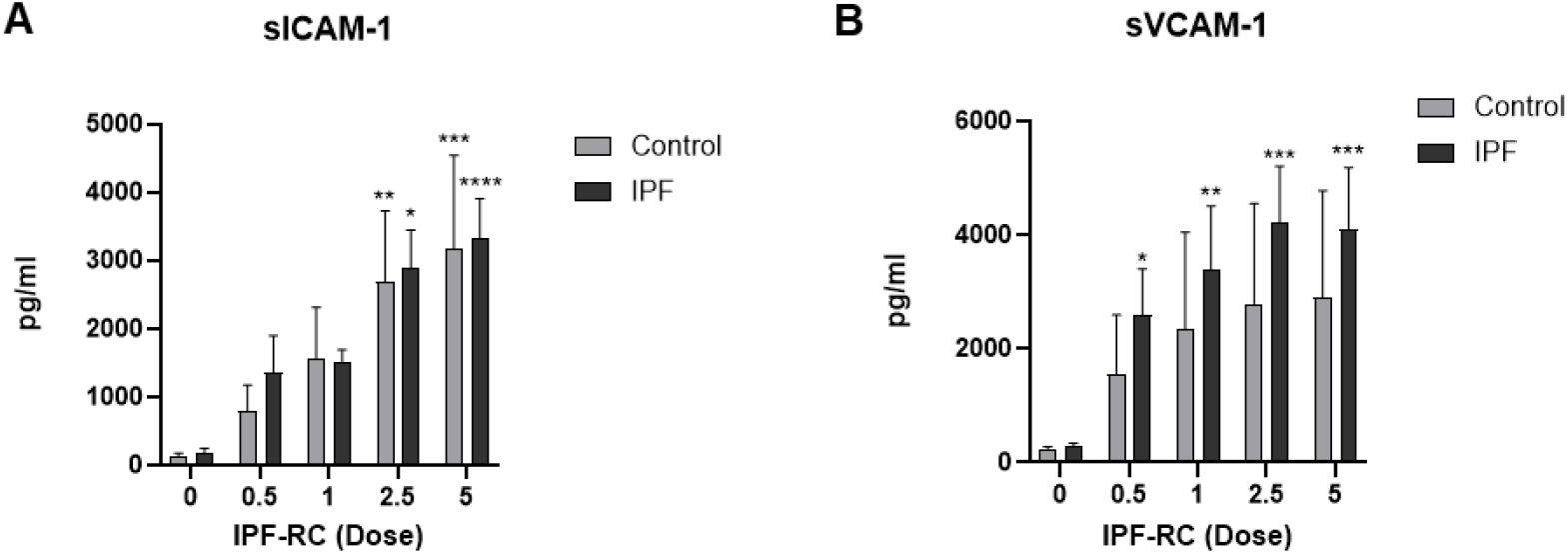
IPF-RC induces dose-dependent endothelial activation in control and IPF-derived ECs. (A–B) ELISA quantification of sICAM1 and sVCAM1 levels in culture media from control (n = 3) and IPF-derived (n = 4) cells treated with IPF-RC for 48 h. Statistics: two-way ANOVA with Tukey’s multiple comparison test. Asterisks indicate comparisons between untreated and treated samples within each group. Data: mean ± SEM. Significance: p ≤ 0.05 (*), p ≤ 0.01 (**), p ≤ 0.001 (***), p ≤ 0.0001 (****). Only significant results are labeled.

Given the plasticity observed in primary microvascular ECs, we extended our analysis to commercially available endothelial cell lines. These included macrovascular Human Umbilical Vein Endothelial Cells (HUVEC), immortalized Human Lung Microvascular Endothelial Cells (HULEC5A), hiPSC-derived ECs, and commercially available primary Human Lung Microvascular Endothelial Cells (HMVEC). This comparison aimed to evaluate their suitability for modeling fibrotic endothelial dysfunction and to identify systems with translational relevance for drug screening.

While the morphology of the commercial endothelial cell lines varied considerably, HMVECs appeared most similar to primary CD31^+^ ECs in terms of visual characteristics (Fig. 5). To assess microvascular identity, we performed PCR analysis comparing these cell types to lung tissue and freshly isolated CD31^+^ ECs. The aerocyte marker HPGD was undetectable in HUVECs, HULEC5a, and HMVECs, whereas primary CD31^+^ ECs and hiPSC-derived ECs retained detectable levels (Fig. 6A, A4). Another aerocyte marker, EDNRB, was present in all cell types except HULEC5a. Overall, HULEC5a cells exhibited the lowest expression of microvascular markers, distinguishing them as the least representative of native lung microvascular ECs. Surprisingly, HMVECs did not outperform HUVECs in terms of microvascular marker expression (Fig. 6A, A3), despite their lung origin.

**Figure 5.**
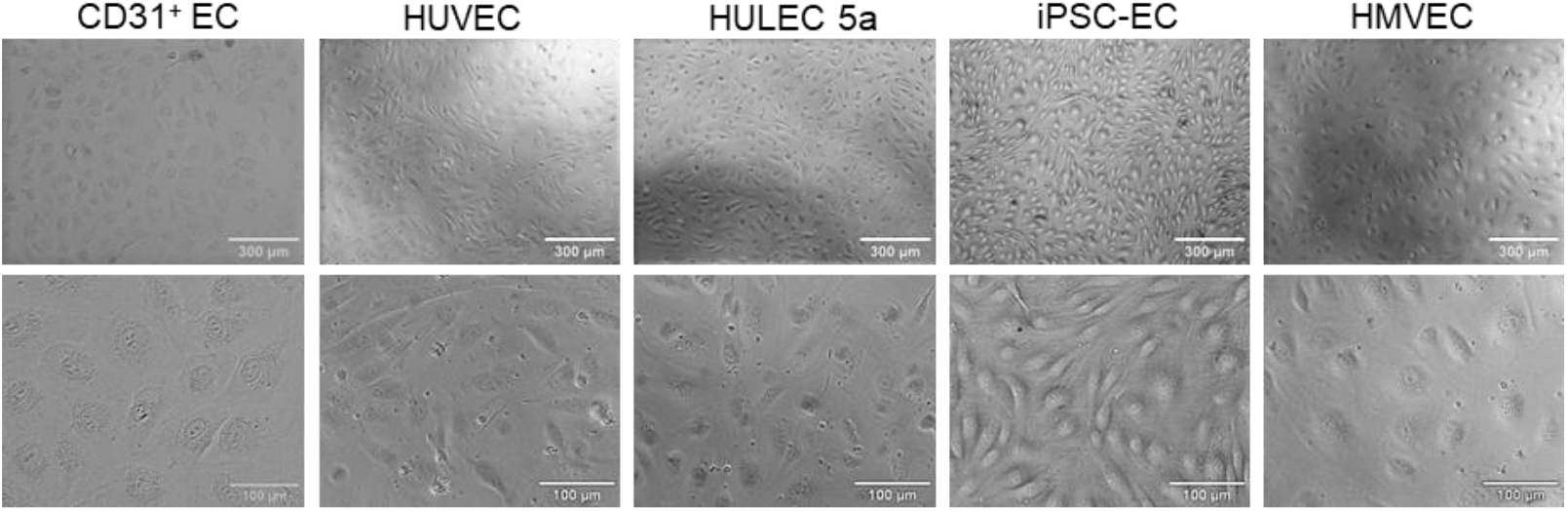
Morphological comparison of primary and commercial endothelial cell lines. Representative brightfield images of endothelial cells (primary CD31^+^ EC, HUVEC, HULEC5a, hiPSC-EC, HMVEC).

**Figure 6.**
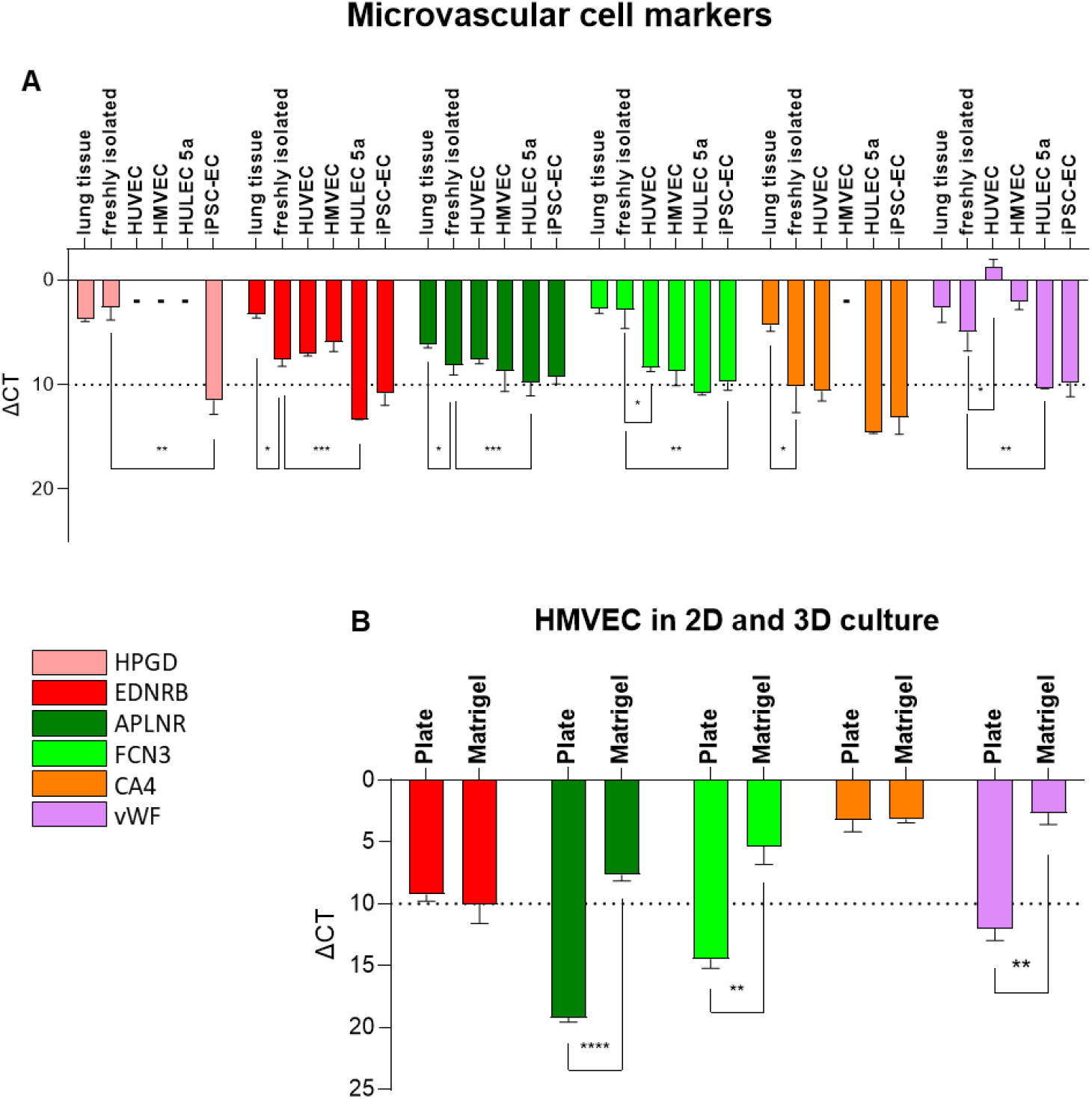
Microvascular identity in ECs: Gene expression analysis and culture condition effects. (A) Expression of microvascular cell markers (ΔCT) in various endothelial cells and lung tissue: HUVEC (n = 4), HMVEC (n = 3), HULEC5a (n = 3), iPSC-EC (n = 3), freshly isolated CD31^+^ ECs (n = 3), and lung tissue (n=5). A minus sign (–) indicates undetectable gene expression. Error bars: SEM from n donors. (B) Expression of endothelial markers in HMVECs cultured on collagen I-coated plates or Matrigel for 8 days. Statistics: two-tailed t-test. Significance: p ≤ 0.05 (*), p ≤ 0.01 (**), p ≤ 0.001 (***), p ≤ 0.0001 (****). Only significant results are labeled; ΔCT > 10 indicates low expression.

Given that HMVECs are primary cells, we hypothesized that the culture environment - specifically the stiffness of plastic substrates - may contribute to dedifferentiation and loss of microvascular identity. To test this, we cultured HMVECs on either standard plastic or soft Matrigel. Notably, Matrigel culture led to increased expression of gCap markers and the ectopic EC marker PLVAP (Fig. 6B), suggesting that endothelial fate is more accurately preserved in physiologically relevant microenvironments.

Despite limitations in marker expression, HMVECs responded to IPF-RC stimulation in a manner comparable to primary CD31^+^ ECs. Upon exposure, they exhibited a dose-dependent increase in inflammatory signaling and a marked reduction in junctional proteins such as VE-cadherin and β-catenin (Fig. 7A–C). Flow cytometry confirmed upregulation of ICAM1 and VCAM1 (Fig. A2B), and ELISA analysis showed elevated levels of soluble ICAM1 and VCAM1 in the culture supernatant. However, we were unable to detect VE-cadherin secretion in monoculture conditions (Fig. 7D), possibly due to intracellular retention or degradation mechanisms.

**Figure 7.**
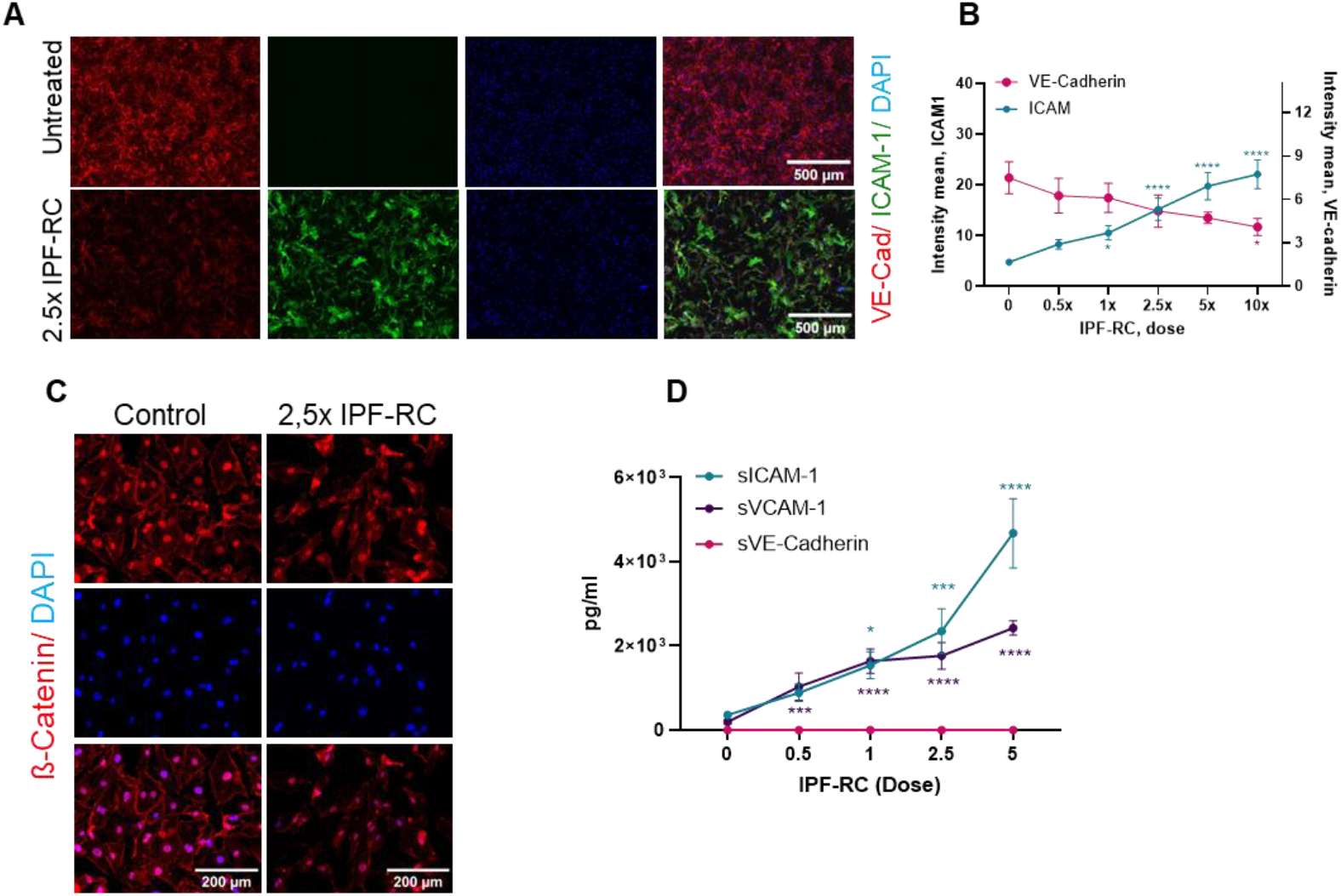
Dose-dependent effects of IPF-RC on junctional proteins and inflammatory markers in HMVECs. (A) IF images of untreated HMVECs and cells treated with 2.5× IPF-RC for 48 h. VE-cadherin (red), ICAM1 (green), DAPI (blue). (B) Quantification of VE-cadherin and ICAM1 IF staining in HMVECs treated with IPF-RC at 0×, 0.5×, 1×, 2.5×, 5× and 10× concentrations for 48 h (n = 3). (C) IF images of β-catenin (red) and DAPI (blue) in HMVECs treated with 2.5× IPF-RC for 48 h compared to untreated cells. (D) ELISA quantification of VE-cadherin, sICAM1, and sVCAM1 in media from HMVECs treated with IPF-RC at 0×, 0.5×, 1×, 2.5×, and 5× for 48 h (n = 3). Statistics: one-way ANOVA with Dunnett’s multiple comparisons. Significance: p ≤ 0.05 (*), p ≤ 0.01 (**), p ≤ 0.001 (***), p ≤ 0.0001 (****). Only significant results are labeled.

Given the well-established role of cyclic AMP (cAMP) in stabilizing endothelial junctions [20,21], we investigated whether a degradation-resistant analog, pCTP-cAMP, could mitigate IPF-RC–induced dysfunction. Treatment with pCTP-cAMP enhanced junctional VE-cadherin expression (Fig. 8A) and partially reversed the deleterious effects of IPF-RC (Fig. 8B). Interestingly, sICAM1 levels remained unchanged in the presence of cAMP (Fig. 8C), suggesting selective rescue of barrier integrity. Functional permeability assays using fluorescent dextrans confirmed that IPF-RC disrupted barrier function, which was fully restored by pCTP-cAMP co-treatment (Fig. 8D, E).

**Figure 8.**
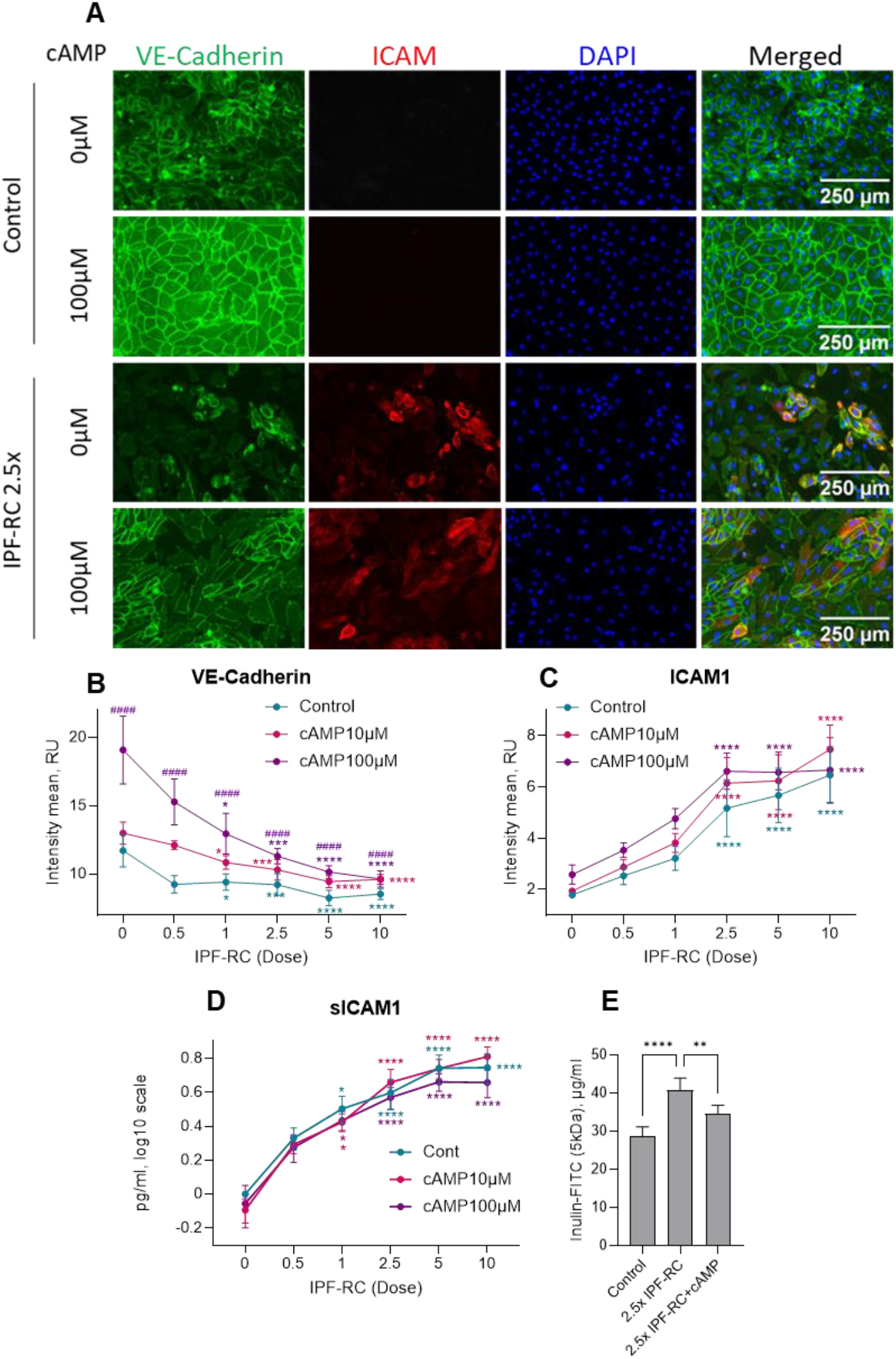
cAMP preserves endothelial barrier integrity under fibrotic conditions. (A) Representative IF images of HMVECs untreated or treated for 48 h with 2.5× IPF-RC ± 100 µM pCTP-cAMP; VE-cadherin (green), ICAM1 (red), DAPI (blue). (B, C) Quantification of VE-cadherin (B) and ICAM1 (C) IF staining in HMVECs treated with IPF-RC at concentrations of 0×, 0.5×, 1×, 2.5×, 5×, and 10×, with or without pCTP-cAMP. Mean intensity per well quantified using ImageJ. (D) ELISA quantification of sICAM1 in media from HMVECs treated with IPF-RC at concentration 0×, 0.5×, 1×, 2.5×, 5×, and 10× for 48 h. Statistics for B, C and D: two-way ANOVA with Turkey’s multiple comparisons test. Error bars: SEM (n = 4). An asterisk (*) indicates a comparison to the untreated control within each group. A hash symbol (#) indicates a comparison between cAMP-treated samples and control samples exposed to the same concentration of IPF-RC. Significance: p ≤ 0.05 (*), p ≤ 0.01 (**), p ≤ 0.001 (***), p ≤ 0.0001 (****, #####). (E) Barrier function assay using Inulin-FITC 5 kDa after 48 h of 2.5× IPF-RC treatment. Statistics: one-way ANOVA with Turkey’s multiple comparisons test. Error bars: SEM (n = 4). Significance: p ≤ 0.05 (*), p ≤ 0.01 (**), p ≤ 0.001 (***), p ≤ 0.0001 (****).

Endothelial dysfunction in IPF is also characterized by impaired cell migration and wound healing [22,23]. To model this, we performed a scratch assay on HMVEC monolayers treated with IPF-RC. Cells exposed to fibrotic stimuli displayed altered morphology and significantly reduced migration capacity (Fig. 9A–C), supporting the hypothesis that IPF-associated cytokines compromise endothelial repair mechanisms.

**Figure 9.**
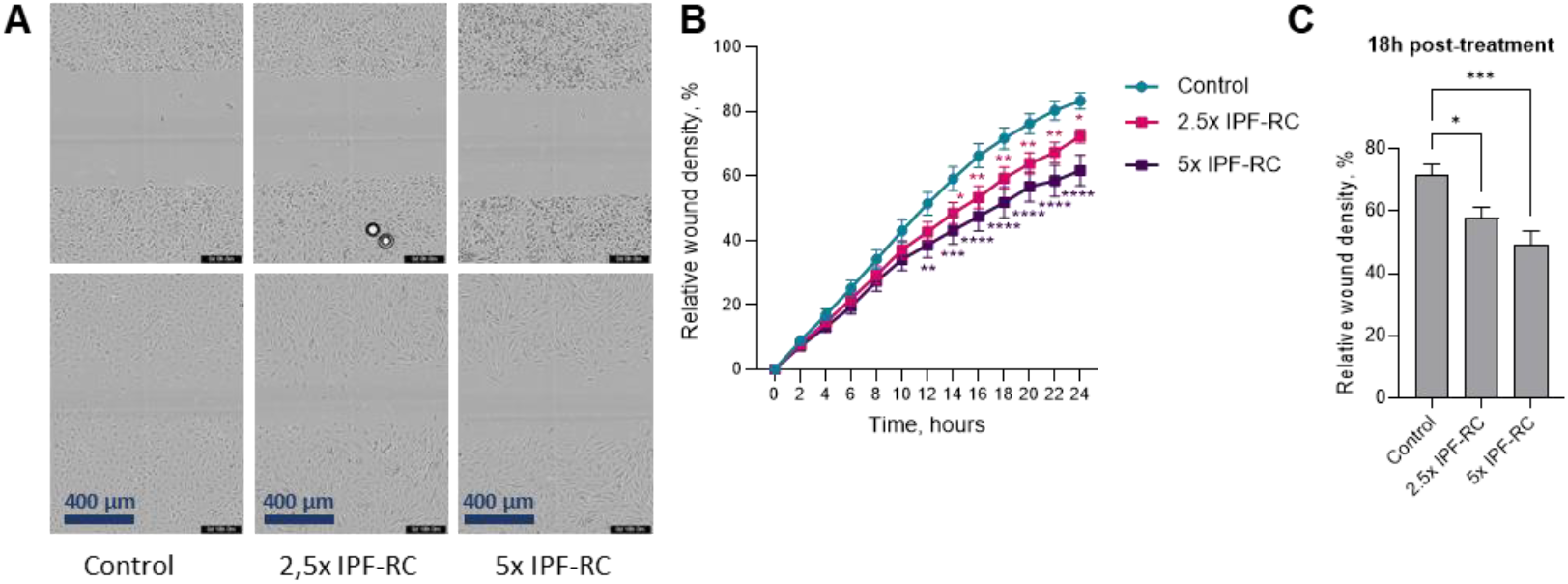
IPF-RC impairs endothelial wound healing capacity. (A) Representative images of HMVECs from the wound healing assay captured at 0 h and 18 h following treatment with IPF-RC at concentrations of 0×, 2.5×, and 5×. Top row: initial scratch; bottom row: cell migration after 18 h. (B) Average relative wound density in the wound healing assay. Statistics: Two-way ANOVA with Dunett’s multiple comparisons test. Error bars: SEM (n = 6). Significance: p ≤ 0.05 (*), p ≤ 0.01 (**), p ≤ 0.001 (***), p ≤ 0.0001 (****). (C) Relative wound density at 18 h. Statistics: one-way ANOVA with Dunett’s multiple comparisons. Error bars: SEM (n=6). Significance: p ≤ 0.05 (*), p ≤ 0.01 (**), p ≤ 0.001 (***).

Another important function of ECs is formation of connections and building of new vessels to repair the injury. Therefore, we used a tube formation assay to study how angiogenic potential of EC is affected in fibrotic environment [24]. We assessed the angiogenic potential of ECs under fibrotic conditions using a tube formation assay. HMVECs treated with IPF-RC formed disrupted networks with fewer connections and reduced structural complexity (Fig. 10A). Quantitative analysis revealed significant reductions in segment number, total branch length, and node count (Fig. 10B–D). Remarkably, co-treatment with pCTP-cAMP restored these parameters to near-control levels, indicating that cAMP not only preserves barrier function but also supports endothelial communication and angiogenesis under fibrotic stress.

**Figure 10.**
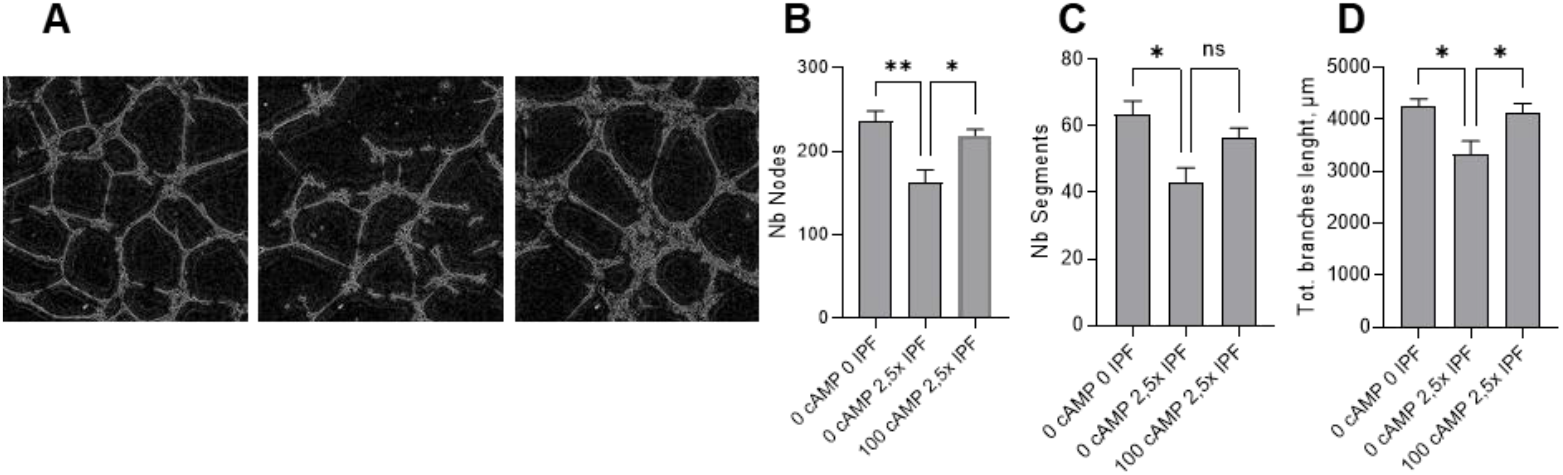
Disrupted Endothelial Tube Formation Under Fibrotic Conditions Is Reversed by cAMP. (A) Representative images of HMVEC cells during a tube formation assay. Cells were seeded on Matrigel, and after 1 hours, the medium was replaced with fresh medium containing 2.5× IPF-RC or 2.5× IPF-RC supplemented with 100 µM pCTP-cAMP. The ability of the cells to form connections was analyzed after 12 hours. (B, C, and D) Quantification of the tube formation assay. The data are represented as the number of segments (B), total branch length (C), and number of nodes (D). Statistics: one-way ANOVA with Tukey’s multiple comparisons. Error bars: SEM (n = 5). Significance: p ≤ 0.05 (*), p ≤ 0.001 (**).

## Discussion

Pulmonary microvascular endothelial cells (PMVECs) are integral to the alveolar niche, contributing to gas exchange, immune regulation, and vascular integrity. Multiple studies have reported a reduction in microvascular ECs in IPF lungs, correlating with fibrotic remodeling and impaired tissue regeneration IPF [7,12,25]. Additionally, reduced endothelial adhesion was found to be accompanied by the formation of fibroblastic foci in usual interstitial pneumonia (UIP) pattern remodeling, as well as fully developed fibrotic areas in alveolar fibroelastosis (AFE) and non-specific interstitial pneumonia (NSIP), as identified by ingenuity pathway analysis of our gene expression results [26]. Our findings support these observations, showing a significant decrease in both aerocyte (aCap) and general capillary (gCap) markers in IPF tissue. These changes likely reflect a loss of specialized EC subtypes, as confirmed by scRNA-seq data from the Lung Endothelial Cell Atlas [11].

Importantly, our study highlights a critical limitation in current endothelial research: the lack of physiologically relevant *in vitro* models. Commercially available ECs, including HUVECs and HMVECs, failed to recapitulate microvascular identity, despite their widespread use. This discrepancy underscores the need for advanced culture systems that preserve endothelial phenotype and function. Similarly, freshly isolated pulmonary CD31^+^ ECs cultured in 2D rapidly lost microvascular identity, eventually resembling commercially available HMVECs. This phenotypic drift likely reflects the limitations of conventional 2D culture systems, which lack critical *in vivo* features, such as unidirectional flow [30], soft basement membranes [31], mechanical stretch from breathing, and niche-specific signaling [32]. To better preserve microvascular characteristics, complex *in vitro* models incorporating those cues are needed [33–36]. A comparable loss of identity has been observed in alveolar type II (AT2) cells and fibroblasts cultured on plastic, where they undergo transdifferentiation and lose their native phenotype [37–39]. Supporting this, culturing HMVECs in Matrigel increased gCap and aCap marker expression, compared to 2D conditions. However, it remains uncertain whether cells can fully regain their original phenotype after 2D expansion. Integrating such advanced culture systems into differentiation protocols for hiPSC-derived ECs may help induce a microvascular phenotype. Although hiPSC-ECs are relatively immature compared to adult cells, they offer higher yields and enable genetic modeling using patient-derived lines or engineered mutations relevant to IPF [40].

Primary cell-based assays – preferentially 2D-models enabling automatic plate readout - remain the gold standard for drug discovery and disease-relevant pharmacology. However, complexity must be broken down by mechanisms to ensure sufficient throughput. To address this, we developed pharmacologically relevant test systems using primary ECs and commercial HMVECs exposed to an IPF-relevant cytokine cocktail (IPF-RC). Notably, the IPF-RC mimics the inflammatory milieu of IPF lavage and offers a more disease-relevant alternative to TGFβ-only stimulation, capturing the complexity of fibrotic signaling [18,19]. IPF-RC induced hallmark features of endothelial dysfunction, including increased ICAM1/VCAM1 expression [41], impaired barrier integrity, and reduced wound healing properties. These responses were consistent across cell types and validated using functional assays. These findings suggest that IPF-related cytokines diminish the regenerative capacity of ECs. This aligns with published data showing disorganized cell migration in fibrotic lungs, which may hinder wound closure [22,23]. Overall, these results underscore the effectiveness of IPF-RC in modeling fibrotic injury in PMVECs.

We also investigated VE-cadherin, known to be downregulated in the plasma of IPF patients [42], but observed no definitive association between its soluble form *in vitro* and fibrotic status in our assays. This lack of correlation may be attributed to injury-induced endocytosis of VE-cadherin [34], potentially mediated by its intracellular interaction with β-catenin, a key component of the Wnt signaling pathway [35]. Immunofluorescence revealed a concurrent decrease in junctional VE-cadherin and β-catenin following IPF-RC treatment. Although further analysis is required, these findings suggest potential activation of the Wnt pathway, which may influence PMVEC fate in IPF lung tissue [43,44].

The translational value of the functional test systems presented here is further demonstrated by our cAMP rescue experiments. cAMP is proposed to enhance barrier integrity [47] and prevent VEGF-induced permeability [48], and phosphodiesterases (PDE)-resistant forms are commonly used to stabilize ECs in culture [20]. In our model, cAMP increased junctional VE-cadherin in untreated HMVECs, indicating improved barrier function. This was confirmed by a dextran-based assay, which showed enhanced barrier integrity in IPF-RC-treated cells. Additionally, cAMP restored EC network formation in tube formation assays. These findings align with clinical data from nerandomilast, a PDE4 inhibitor, which showed positive outcomes in Phase III trials for IPF and PPF, with strong responses in HMVECs [49–51]. Since PDE4 degrades cAMP, using degradation-resistant cAMP might mimic PDE4 inhibition. Similar effects were observed with roflumilast, another PDE4 inhibitor approved for COPD [52]. Together, these results suggest that maintaining cAMP levels may offer therapeutic benefits by protecting the microvasculature.

Moreover, our results and the literature suggest that endothelial activation is not merely a bystander effect but a potential driver of fibrosis. The observed changes in morphology, migration, and barrier integrity suggest that ECs contribute to the perpetuation of injury and impaired repair. This reinforces the need for test systems that can dissect EC-specific mechanisms and evaluate therapeutic interventions in a controlled, reproducible manner.

In conclusion, our study provides compelling evidence that PMVECs are highly responsive to fibrotic stimuli and may play an active role in IPF pathogenesis. By establishing robust pharmacological test systems, we offer a platform for mechanistic studies and drug screening aimed at restoring endothelial function. These models bridge the gap between basic research and clinical translation, supporting the development of targeted therapies for IPF and related fibrotic lung diseases.

## 4. Materials and Methods

### 4.1. Patient samples

Control and fibrotic human distal lung tissue samples were obtained from patients at Hannover Medical School (MHH), with written informed consent provided for research purposes. The study received approval from the local ethics committee at MHH (8867_BO_K_2020) and was conducted in accordance with their guidelines. The IPF cohort comprised six male patients aged 58 to 68 years, all presenting with clinically progressive fibrosis. The diagnosis was confirmed through radiological and histopathological assessment, which identified usual interstitial pneumonia (UIP) patterns. For the control group, healthy lung tissue was sourced from five donors (two males and three females aged 54 to 72 years old) who had undergone lung tumor resections. The lack of macroscopic and microscopic alterations was confirmed by two experienced pulmonary pathologists. Neither the patients nor their families received any compensation for the IPF patient samples or the control lung tissues.

### 4.2. Cell isolation from human lung tissue

Human lung tissue was minced with scissors in a petri dish and then transferred to tubes containing DMEM medium (5 ml per 1 g tissue) along with digestive enzymes from the tumor dissociation kit from Miltenyi (130-095-929). The tissue was incubated at 37°C for 60 min using the gentleMACS™. To inactivate the digestion, 1 ml of FBS was added, and the cell suspension was filtered through 500 µm, 300 µm, and 100 µm strainers to remove debris. The cells were then centrifuged at 300 × g for 10 minutes at 4°C, and an erythrocyte depletion buffer (420301 Biolegend) was applied according to the manufacturer’s instructions. The cells were filtered again using a 40 µm strainer, washed, and centrifuged. The prepared cells were resuspended in MACS Separation Buffer (MILT130-091-376, Miltenyi Biotec + 5% MACS® BSA Stock Solution 130-091-376) and blocked with FcR Blocking Reagent (MILT130-059-901, Miltenyi Biotec) for 3 min. In the first step of cell sorting, CD45-positive cells were removed using CD45 MicroBeads (MILT130-045-801, Miltenyi Biotec). After washing and centrifugation, CD31-positive cells were collected using CD31 MicroBeads (MILT130-091-935, Miltenyi Biotec). For expansion, the sorted cells were seeded on collagen I-coated plates and supplemented with EGM2 MV media (CC-3202, Lonza) with 20% FBS and 10µM Rock-Inhibitor (SCM075, Millipore). Rock-Inhibitor was removed from the culture media after three days. Expanded cells were sorted via FACS using anti-CD31 antibodies on a Sony MA900 Cell Sorter to isolate CD31^+^ populations with high purity. After the second sorting, the cells were cultured in EGM2 MV media with 5% FBS.

### 4.3. Commercially available endothelial cells

Human umbilical vein endothelial cells (HUVEC, C-12200) and human lung microvascular endothelial cells (HMVEC-L, CC-2527) were purchased from PromoCell and Lonza, respectively. Human lung microvascular endothelial cells (HULEC-5a, CRL-3244) were obtained from ATCC (Manassas, USA). All assays were performed with cells at passages 2-6.

### 4.4. Differentiation of hiPSC derived endothelial cells

Human induced pluripotent stem cell-derived endothelial cells (hiPSC-ECs) were differentiated according to previously published protocols [53,54], with minor adaptations. In summary, 30-40 colonies of hiPSCs per well were seeded in mTeSR Plus medium (C-100-0276, STEMCELL TECHNOLOGIES) on GFR Matrigel (Corning)-coated six-well plates (C-140675, Thermo Scientific). Mesoderm induction was initiated 24 hours later by switching the medium to bovine serum albumin (BSA) polyvinyl alcohol essential lipids (B(P)EL) medium, supplemented with 8 µM CHIR99021 (C-4423/10, Tocris Bioscience). Vascular specification was induced on day 3 by transitioning to B(P)EL medium supplemented with 50 ng/ml VEGF (MILT130109384, Miltenyi Biotec) and 10 µM SB431542 (C-1614, Tocris Bioscience). Media was refreshed on days 6 and 9. To enrich the purity of endothelial cells, CD31-Dynabeads were used (11155D, Thermo Fisher), after which the cells were further expanded for 4 days. The hiPSC-ECs were used in passage 2.

### 4.5. PCLS

Explanted lungs were infused with 1.5% low-gelling temperature agarose and incubated on ice until solidified. Tissue punches (5 mm) were generated using a tissue coring press. Precision-cut lung slices (PCLS) with a thickness of 200-300 µm were prepared using a Krumdieck tissue slicer and were immediately transferred to Netwell™ insert plates containing CPC media, where they were gently shaken in an incubator set to 37°C. To reduce the agarose content, the PCLS were washed for 30 minutes six times, followed by a final overnight wash. The next day, the PCLS were transferred to 48-well plates, one slice per well in 300 µL of CPC medium, and treated with various concentrations of TGFβ. After 48 hours of treatment, samples were collected for RNA analysis to evaluate the effects of TGFβ. Three PCLS per condition were obtained from a total of three individual donors.

### 4.6. Wound healing assay

HMVEC cells were seeded onto a 96-well plate (Sartorius, BA-04855) at a density of 20,000 cells per well and incubated in a culture medium supplemented with 5% FCS until a confluent monolayer was formed. An artificial wound was created using an automated 96-pin wound-making tool (WoundMaker™; Essen Bioscience). The medium was then replaced with a medium containing 1.5% FCS and varying concentrations of IPF-RC [14]. Cells were maintained for 24 hours in an IncuCyte™ device (Sartorius). Images were captured every two hours. Relative wound density was calculated using the IncuCyte™ software.

### 4.7. Tube formation assay

A 96-well plate was coated with 50 µl of Matrigel (356234, Corning) overnight. HMVECs, were seeded onto the prepared plates at a density of 10,000 cells per well. After an incubation period of 1 hours, the media was replaced with one of the following: no compounds, 2.5x IPF-RC, or 2.5x IPF-RC supplemented with 100µM pCTP-cAMP. The tube formation process was then observed and imaged at the 12-hour time point using the Opera Phenix® Plus High-Content Screening System. The images were analyzed using the Fiji Plugin Angiogenesis Analyzer.

### 4.8. Barrier function assay

Inserts for 24-well plates with 0.4 µm pores size were coated with collagen I and incubated for 1 hour. Following incubation, the coating solution was removed, and 20,000 cells were seeded onto each insert. After 24 hours, cells were starved for 1.5h with a medium containing 1.5 % FBS, then treated with 2.5x IPF-RC for 48 hours. To assess barrier integrity, a permeability assay was performed using Inulin-FITC (5 kDa), diluted to 1 mg/mL in phenol red free EGM medium (CC-3129, Lonza). A volume of 100 µL of the FITC-labeled solution was added to the apical compartment of each insert. After 2 hours of incubation, media from the basolateral compartment was collected, and fluorescence intensity was measured using a SpectraMax M5 spectrophotometer (Molecular Devices) with excitation at 495 nm and emission at 519 nm. Data were quantified using a standard curve.

### 4.9. Immunofluorescence

Cells were fixed with 4% paraformaldehyde for 30 minutes at room temperature, followed by washing with phosphate-buffered saline (PBS). Permeabilization was performed using 0.1% Triton X-100 (CAS 9036-19-5, Sigma) for 20 minutes. Cells were then blocked with 5% donkey serum for 1 hour at room temperature. Primary antibody staining was performed overnight at 4°C using the following antibodies: anti-ICAM1 Alexa 647 (ab214944, Abcam), anti-VE-cadherin PE (12144982, Invitrogen), and anti-β-Catenin (05-665-25UG, Merck). For β-Catenin detection, cells were incubated with a secondary antibody—donkey anti-mouse Alexa Fluor 647 (A11015, Invitrogen)—for 1 hour at room temperature. Nuclear staining was performed using DAPI (62248, Thermo Fisher Scientific) for 5 minutes at room temperature. Fluorescence images were acquired using the EVOS M7000 microscope (Thermo Fisher Scientific) and analyzed with Fiji (ImageJ) software. A complete list of antibodies used is provided in Supplement 1, Table 1.

### 4.10. Real time PCR

Total mRNA was isolated using either the RNeasy Plus Micro Kit (Qiagen, Irvine, CA, USA) or the Maxwell RSC Simply RNA Kit (Promega). The concentration and purity of the mRNA were then analyzed using a Nanodrop spectrophotometer. cDNA was synthesized with the High-Capacity cDNA Reverse Transcription Kit (Applied Biosystems). Quantitative PCR (qPCR) was conducted and analyzed using the ViiA 7 Real-Time PCR System (Applied Biosystems). A list of the primers used can be found in Appendix 1, Table 2. For each sample, qPCR was performed in triplicate with both a gene of interest primer and a housekeeping primer (ACTB).

### 4.11. ELISA

Enzyme-linked immunosorbent assays (ELISA) were conducted following the manufacturer’s instructions for the following proteins: ICAM1 (DY720, Biotechne R&D Systems), VE-cadherin (DY938-05, Biotechne R&D Systems), and VCAM1 (DY809, Biotechne R&D Systems). Signal detection was performed using the SpectraMax M5 instrument from Molecular Devices.

### 4.12. Flow cytometry and FACS

Single-cell suspensions were first stained with Zombie dye (423107, BioLegend) at a 1:400 dilution before fixation. Next, the eBioscience™ Foxp3/Transcription Factor Staining Buffer Kit (00552300, Invitrogen) was used for fixation and staining, following the manufacturer’s instructions. Cells were then incubated with primary antibodies for 30 minutes at room temperature, followed by a 30-minute incubation with secondary antibodies (see Appendix 1 for the list of antibodies used). For live cell staining, FACS buffer was utilized as described by Wasnick et al. **[55]**. Nuclear counterstaining was performed using DAPI (62248, ThermoFisher Scientific) at a dilution of 1:2000. Flow cytometry was conducted using the BD LSRFortessa X-20 (BD Biosciences), and cell sorting was carried out with the SONY MA900 Multi-Application Cell Sorter. Finally, data analysis was performed using FlowJo version 10.10.0 software (Tree Star Inc., Ashland, OR, USA).

### 4.13. Statistical analysis

Statistical analyses were performed using GraphPad Prism 10 software. Data are presented as the standard error of the mean (SEM). A Student’s t-test was used for comparisons between the two groups. An ordinary one-way ANOVA with Dunnett’s multiple comparisons was employed to compare several groups to the control. Ordinary one-way ANOVA with Tukey’s multiple comparisons test was used for comparisons among several groups. Additionally, an ordinary two-way ANOVA with Tukey’s multiple comparison test was utilized to compare several groups under various conditions.

### 4.14. Human single cell mRNA sequencing data analysis

Endothelial cell-specific target genes were selected and evaluated based on an integrated dataset of human scRNA-seq data from control and IPF patients, as previously published [8]. In brief, publicly available scRNA-seq datasets of interstitial lung disease (ILD) were integrated into a comprehensive atlas of PF-ILD patients, available on Zenodo (10.5281/zenodo.10015169). For this study, the data was subset to include only cells from IPF and control patients and further refined to include only cells from the endothelial compartment, resulting in a dataset of 12,386 cells. The fold change between IPF and control was calculated using Scanpy’s (v1.10.4) rank_genes_groups function **[56]**.

## Acknowledgments

We thank the BIU3.0 collaboration program from Boehringer Ingelheim and Ulm university for funding and the full I&R team for support. We thank the team of AG Lung research for excellent technical support and Holger Schlueter for coordinating lung deliveries. Tarek Gensheimer expresses gratitude to his supervisor, Stefan Kreideweiss, and acknowledges the support of his PostDoc grant, awarded through the opn2TALENTS call on opnMe.com.

## Institutional Review Board Statement

The study was conducted in accordance with the Declaration of Helsinki and approved by the local ethics committee at MHH (protocol code 8867_BO_K_2020 and approved on 28 January 2020).

## Informed Consent Statement

Control and fibrotic human distal lung tissue samples were obtained from patients at Hannover Medical School (MHH), with written informed consent provided for research purposes.

## Appendix A

**Table A1.** Primary antibodies.

| Target | Dilution | Spec. | company | Cat.number | conjugate |
| --- | --- | --- | --- | --- | --- |
| ICAM-1 | 1:100 | mus | abcam | AB214944 | Alexa 647 |
| VE-Catherin | 1:100 | mus | Invitrogen | 12144982 | PE |
| P16<sup>INK4a</sup> | 1:100 | mus | Invitrogen | MA5-17054 | - |
| β-Catenin | 1:100 | mus | Merck | 05-665-25UG | - |

**Table A2.**
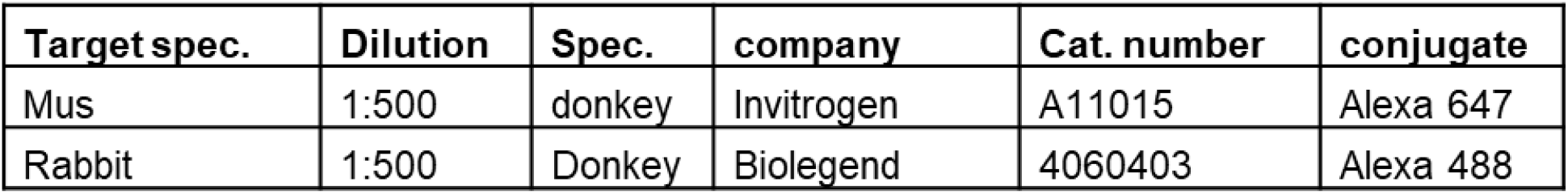
Secondary antibodies.

**Table A3.** Compounds.

| Substance | Concentration | Source |
| --- | --- | --- |
| Collagen-I | 5 µg / cm<sup>3</sup> | 354249, Corning |
| Collagen-IV | 200 µg / mL | C5533, sigma |
| Laminin | 5 µg / mL | 521LN-05, Biolamina |
| Fibronectin | 30 µg / mL | 356008, Corning |
| Gelatin | 0.1% | 6950, Cellbiologics |
| pCPT-cAMP | 0,25 mg/ml | C3912, Sigma Aldrich |

**Table A4.** Ingredients of 1000x IPF-RC.

| component | Concentration | Source |
| --- | --- | --- |
| TGF-β1 | 0.30 µg / mL | 240-B-002, R&D |
| IL-1β | 0.01 µg / mL | 201-LB-005, R&D |
| TNF-α | 0.10 µg / mL | 210-TA-020, R&D |
| IL-8 | 1.50 µg / mL | 208-IL-010, R&D |
| MCP1 (CCL2) | 0.70 µg / mL | 279-MC-010, R&D |
| IL-33 | 0.04 µg / mL | 3625-IL-010, R&D |
| TSLP | 0.10 µg / mL | 1398-TS-010, R&D |
| IL-13 | 2.50 µg / mL | 213-ILB-025, R&D |
| IL-4 | 0.16 µg / mL | 204-IL-010, R&D |

## Appendix B

**Figure A1.**
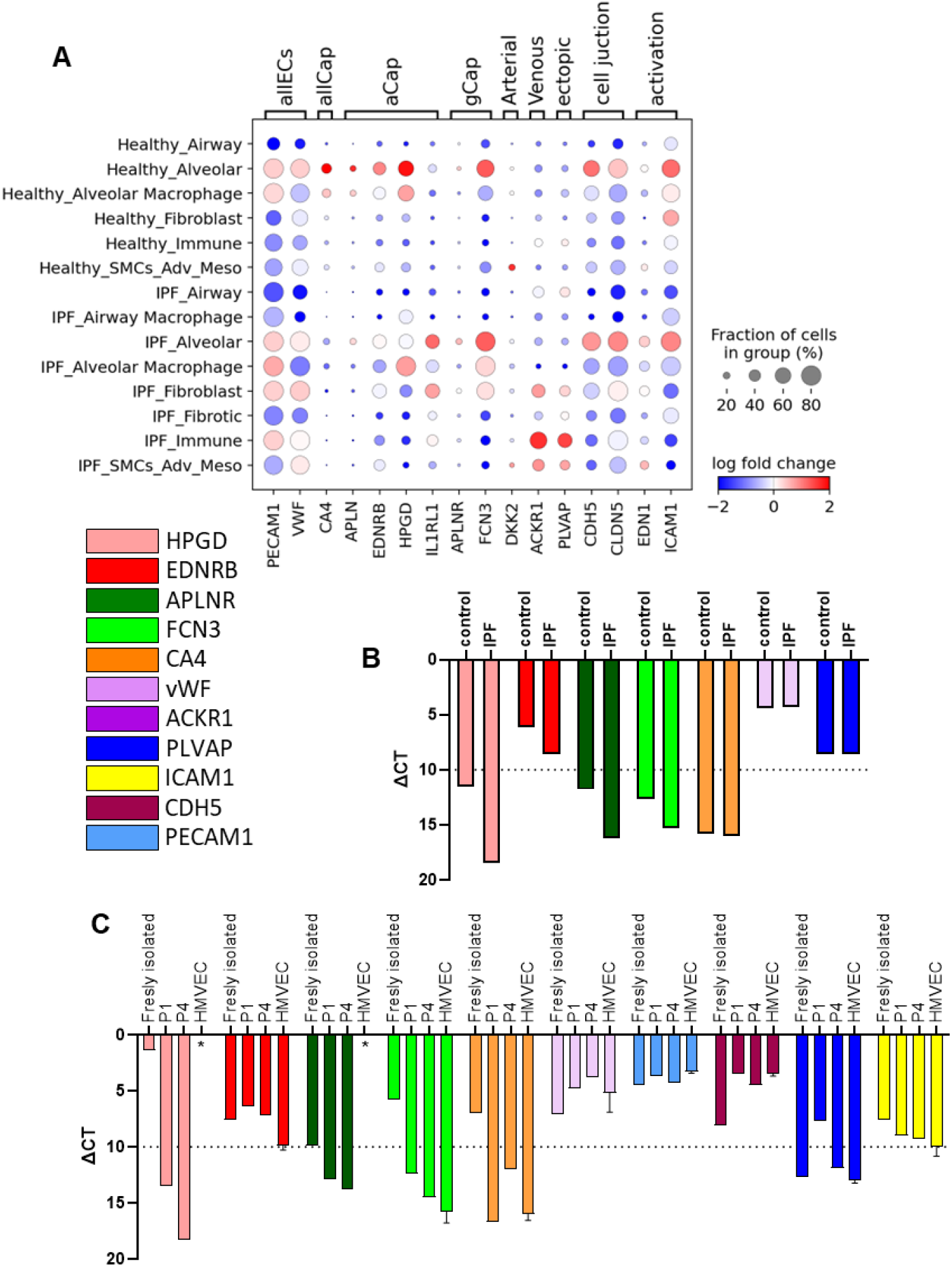
**A** Dot plots illustrating the fold change in expression of indicated marker genes between IPF and control groups across various cell types in scRNA-seq data from IPF and control patients. Data set from Mayr et al., 2024. **B** Expression of microvascular cell markers in freshly isolated CD31^+^ EC from control and IPF lung tissue (n=1), expressed as ΔCT. **C** Expression of microvascular cell markers in freshly isolated and at passage 1 and 4 CD31^+^ EC, expressed as ΔCT. Data from HMVEC cells are provided for comparison. Asterisks (*) indicate undetectable levels of gene expression. Error bars represent the standard error of the mean, n=2.

**Figure A2.**
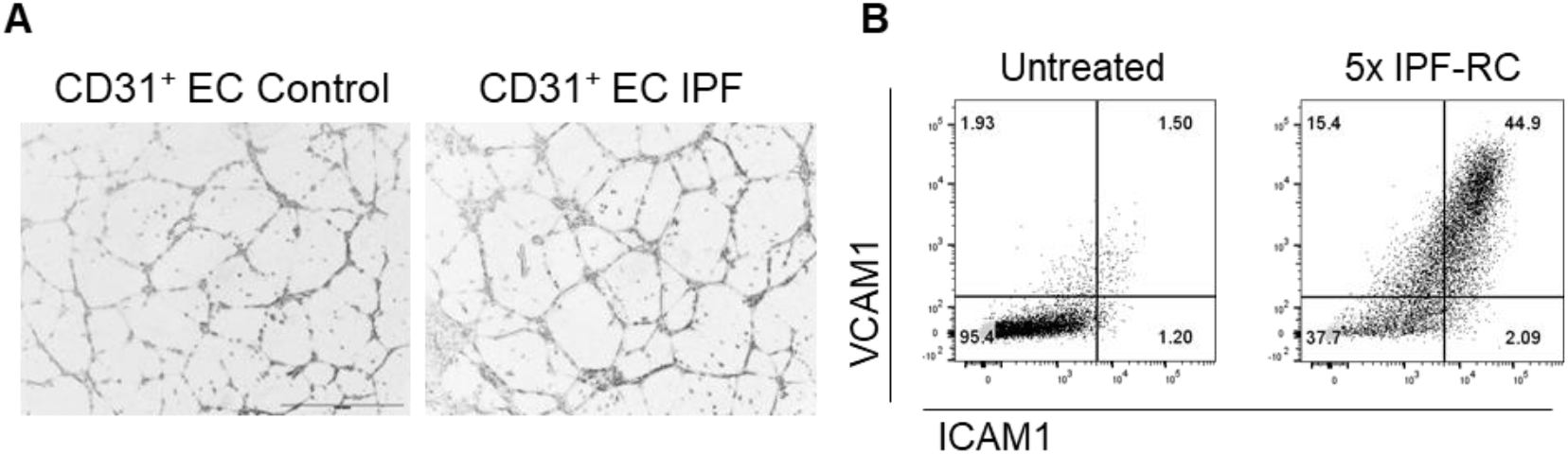
**A** Representative brightfield images of the tube formation assay for primary CD31^+^ endothelial cells (ECs) from idiopathic pulmonary fibrosis (IPF) and control lung tissue. **B** Dot plots from flow cytometry analysis of HMVEC cells treated with IPF-RC for 48 hours, followed by staining with anti-ICAM1 and anti-VCAM1 antibodies.

**Figure A3.**
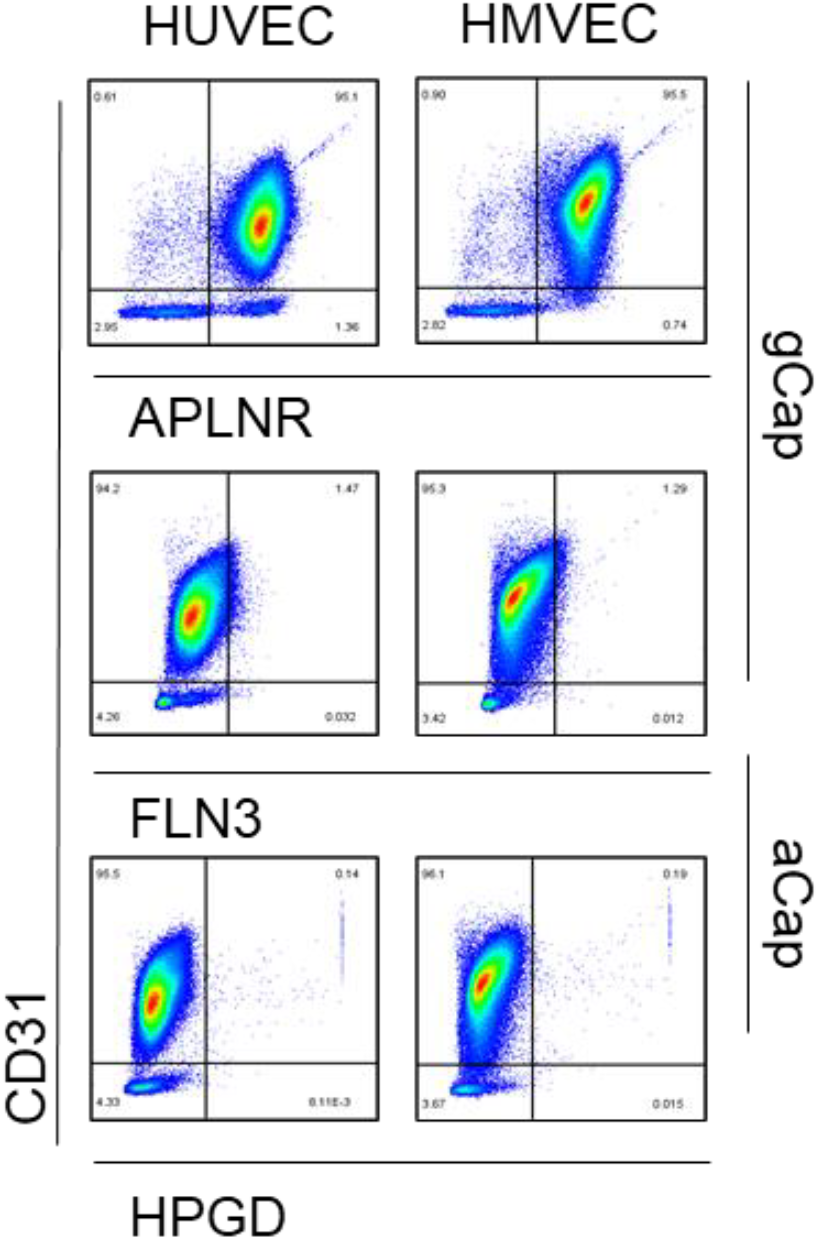
Dot plots from flow cytometry analysis of HUVEC and HMVEC cells with anti-CD31, HPGD, FLN3, and APLNR antibodies.

